# Parental rejection is associated with extended lifespan in owl monkeys in captivity

**DOI:** 10.64898/2025.12.18.695178

**Authors:** Jayde Farinha, Nofre Sánchez-Perea, Ping Yip, Ursula M. Paredes

## Abstract

Parental rejection of apparently healthy newborns is widely classified as a behavioural abnormality in captive primate colonies, yet its biological significance remains unclear. In owl monkeys (*Aotus nancymaae*), parental rejection, defined here as cessation of nursing leading to rescue nursery rearing, is typically lethal for offspring and is transmitted across generations despite reducing offspring survival. Here, we tested whether parental rejection is associated with lifespan and reproductive differences in parents and their surviving offspring. We analysed long-term demographic records from a captive colony of 962 individuals and compared survival and reproductive outcomes between rejector and non-rejector parents using survival analyses and regression-based models.

Parents who rejected offspring lived significantly longer than non-rejectors, with an average lifespan advantage of approximately 4-4.5 years in both males and females. This survival difference was concentrated during the prime reproductive period (6-20 years). Well-reared offspring of rejector parents also lived longer than offspring of non-rejectors, with a mean lifespan difference of 1.26 years. Rejector parents produced more offspring overall, but this difference was explained by extended lifespan rather than higher reproductive output per year. Analyses stratified by rejection timing showed no longevity advantage in first-birth rejectors, whereas parents rejecting later-born offspring exhibited longer survival.

Together, these findings show that parental rejection is associated with longer lifespan in parents and in their well-reared offspring under captive conditions. These patterns are consistent with altered allocation of parental investment under energetic or environmental stress.

## Introduction

Parental rejection of newborns, defined as the cessation of nursing and abandonment of milk-dependent infants, is a widespread phenomenon observed across captive primate colonies, including owl monkeys (*Aotus nancymaae*)^1^. In the absence of assisted rearing, this behaviour is typically lethal. Our follow-up studies reveal wide-ranging effects on survivors, including poorer health outcomes^2^, accelerated epigenetic ageing in blood^3^, intergenerational transmission of reduced odds of survival to reproductive age, shortened lifespans, and altered stress behaviours^4^. More recently, we demonstrated that within the rejected cohort, the experience-induced tendency to reject newborns is transmitted intergenerationally and associated with alterations in epigenetic markers in naïve offspring^5^.

Complete rejection and abandonment of newborns is not unique to this colony of owl monkeys. It has been documented across wild primate populations, including New World monkeys such as tamarins^6^, marmosets^7,8^ and other owl monkey species (Fernandez-Duque, personal communication, January 2024), Old World monkeys^9-11^, and humans^12^, suggesting that the capacity for abandonment represents a conserved, albeit extreme, behavioural strategy. In captivity such behaviours are reported at substantially higher frequencies across multiple primate taxa, including other owl monkey colonies^13-17^, and in humans exposed to stressful environments^12^, indicating that environmental sensitivity may trigger their expression. Importantly, intergenerational transmission has been widely documented for maternal rejecting tendencies more broadly, including normative or experience-dependent reductions in care, in humans^18^, macaques^9, 19-22^ and baboons^23^. In contrast, evidence for the intergenerational transmission of complete neonatal abandonment remains rare; in owl monkeys, we have previously demonstrated such transmission associated with early-life experience^5^.

In owl monkeys, rejection in captivity typically involves maternal refusal to nurse leading to abandonment, and differs fundamentally from normal weaning, in which females gradually transfer care of the young to fathers while continuing to nurse until weaning^24^. A substantial body of literature links offspring neglect to maternal stress during pregnancy, early caregiving experience, and environmental stressors across the lifespan^25-27^. However, why such a severe form of rejection, with profound intergenerational negative consequences for offspring fitness, would nonetheless be transmitted across generations remains unresolved.

Genetic propensity for neglect has been considered. Stress reactivity and gene variants regulating catecholamine neurotransmission are proposed to moderate effects of negative early life^28^ and have been associated with manifestation of maternal neglect and rejecting tendencies^29,30^. Studies of rhesus macaques in captivity suggest that stressed mothers or those in poor condition exhibit higher rejection rates^31^. Rejecting newborns is considered pathological^13^, although, some have proposed that when it occurs in first births, it rather reflects lack of experience^14,15, 27, 32^. As captivity itself can induce chronic stress in some primates^33^, rejecting might be intensified in these environments.

It has also been proposed that offspring rejection and abandonment during the weaning period could be potentially beneficial to mothers. Under natural weaning conditions, both high-ranking and low-ranking vervet monkey mothers exhibit increased rates of maternal rejection compared to average-ranking mothers^11^. The authors proposed that this U-shaped pattern may reflect different strategies: low-ranking mothers conserve energy for self-preservation, while high-ranking mothers accelerate weaning to reproduce again sooner.

More extreme manifestations of rejection have also been suggested as adaptive. In wild marmosets, infant abandonment occurs when rearing conditions are suboptimal, for example, subordinate females’ offspring may be expelled from the group when helpers are scarce^7^. For rhesus macaques in captivity, neonatal abandonment and neglect have been interpreted as reflecting a reduction in parental investment under suboptimal conditions, and, in its most extreme forms, may represent an adaptive response to constrained maternal condition^31,34,35^. In contrast, infant physical abuse appears to be a maladaptive form of aggression rather than a strategic reduction in care^31^. Studies in wild hanuman langurs document instances when, a new male took over a group, he often killed infants he did not sire. Mothers sometimes abandoned or stopped defending their babies to survive and reproduce again sooner^36^. In extreme cases, abandonment followed by infanticide and cannibalism in new world monkeys might reflect maternal nutritional stress^6,8^.

If adaptive, cessation of newborn care has been explained as parent–offspring conflict^37^. Under duress, offspring demands for care and feeding come into conflict with mothers’ needs to survive, repair somatic damage, and invest in future reproduction. When maternal costs of care outweigh benefits, some have argued that selection may favour infant rejection^38^. Further, it has been proposed that the adaptive value of maternal investment to offspring must be weighed on maternal lifetime fitness rather than offspring survival alone^39^. Reduction of maternal investment, through rejection, may restore energy needed for somatic repair, leading to gains in lifespan (Disposable soma theory^40^). In owl monkeys, which reproduce annually^41^, reproductive output is closely tied to lifespan, as observed elsewhere^42^. Thus, maternal extended survival should directly translate to increased reproductive output. In the case of owl monkeys, which uniquely displays biparental care^43^, shared investment may influence how both sexes experience the costs and possible benefits of rejecting behaviour.

In this colony, we have identified a link between rejecting behaviour and expected lifespan and ageing correlates. Rejected owl monkeys which survive have shorter lives and suffer poorer health than controls and suffer from lower biological fitness, effects which are intergenerationally transmitted^4^. Rejection is also associated with accelerated epigenetic ageing in blood^3^. Simultaneously, we previously detected that some mother rejectors enjoy long lifespans^5^. These findings suggest possible links between rejecting behaviour, fitness and survival, although these relationships are unclear.

To investigate the rejecting paradox, we test whether rejecting newborns is associated with increased survival and reproductive output in mothers or both parents, and whether their non-rejected offspring also show enhanced survival and reproductive outcomes.

## Methods

### Study population and housing

This study reanalysed long-term demographic and behavioural records from colonies maintained at the Centre for Reproduction and Conservation of Non-Human Primates (CRCP) of IVITA at the National University of San Marcos (UNMSM), located in Iquitos, Peru. Animals were housed in standardised single enclosures (2 m^3^) with access to natural light cycles and were provided with a diet of seasonal fruits, balanced in house dry cookies, and water *ad libitum*.

### Definition of rejection

Parental rejection was operationally defined as maternal cessation of nursing, resulting in infant malnutrition or lethality risk and necessitating nursery rearing. In owl monkeys, fathers assume primary infant-carrying duties within days of birth but cannot compensate for maternal nursing cessation. Rejected infants were removed for hand-rearing when mothers discontinued nursing. Occasionally, fathers displayed aggressive behaviour (e.g., biting) toward non-nursed infants, but this was rare and secondary to maternal nursing cessation. A total of 26 rejected individuals were identified, compared with 729 non-rejected controls. There were 43 rejectors (Females = 19, Males =24) compared with the control group of 236 non-rejecting parents (Females = 112, Males = 124).

An overview of the datasets and sample sizes used across analyses is provided in Table 1. Importantly, the form of rejection studied here, characterised by neonatal cessation of nursing leading to rescue nursery rearing, differs fundamentally from the ‘rejection’ described in much of the primate literature, which often refers to normative or accelerated weaning behaviours^11,27^ occurring later in infancy and not associated with offspring lethality^1,10^.

**Table 1.** Summary of datasets and sample sizes used in analyses.

| Analysis / Figure | Dataset description | Rejectors (F/M) | Non-rejectors (F/M) | Total N |
| --- | --- | --- | --- | --- |
| Parental lifespan (Fig. 1a, 2a) | All parents with known age at death | 19 / 24 | 112 / 124 | 279 |
| Offspring lifespan (Fig. 1b) | Non-rejected offspring only | 125 / 106 | 253 / 271 | 755 |
| Rejection timing (Fig. 2b) | Parents with known rejection timing | 5+22+11<sup>†</sup> (subset of above) | Included as reference | 274<sup>*</sup> |
| Later birth/repeat rejector vs comparison group<sup>‡</sup> | Aggregated parent groups | 27 | 247 | 274 |
| Reproductive output (Fig. 3) | Parents with lifetime reproductive histories | 43 | 236 | 279 |

<sup>\*</sup> Subset excludes rejectors for whom the rejected birth could not be determined. <sup>†</sup> Repeat rejectors (n = 5), single non-first rejectors (n = 22), and first-birth rejectors (n = 11). <sup>‡</sup> Later-birth/repeat rejectors include individuals rejecting offspring beyond the first birth (single non-first and repeat rejectors); comparison group includes first-birth rejectors and non-rejectors.

### Parents Survival analyses

We investigated lifespan differences between rejector and control parents, and between offspring with or without rejected siblings. For parental analyses, we compared age at death between rejecting mothers (n = 19) and non-rejecting control mothers (n = 112), and between rejecting fathers (n = 24) and non-rejecting control fathers (n = 124). Owl monkeys form monogamous pairs, and rejection events were accounted for both parents. The slight discrepancy in sample sizes reflects mortality and unpaired individuals at the time of rejection. Welch’s two-sample t-tests were used for these comparisons (Figure 1a).

**Figure 1.**
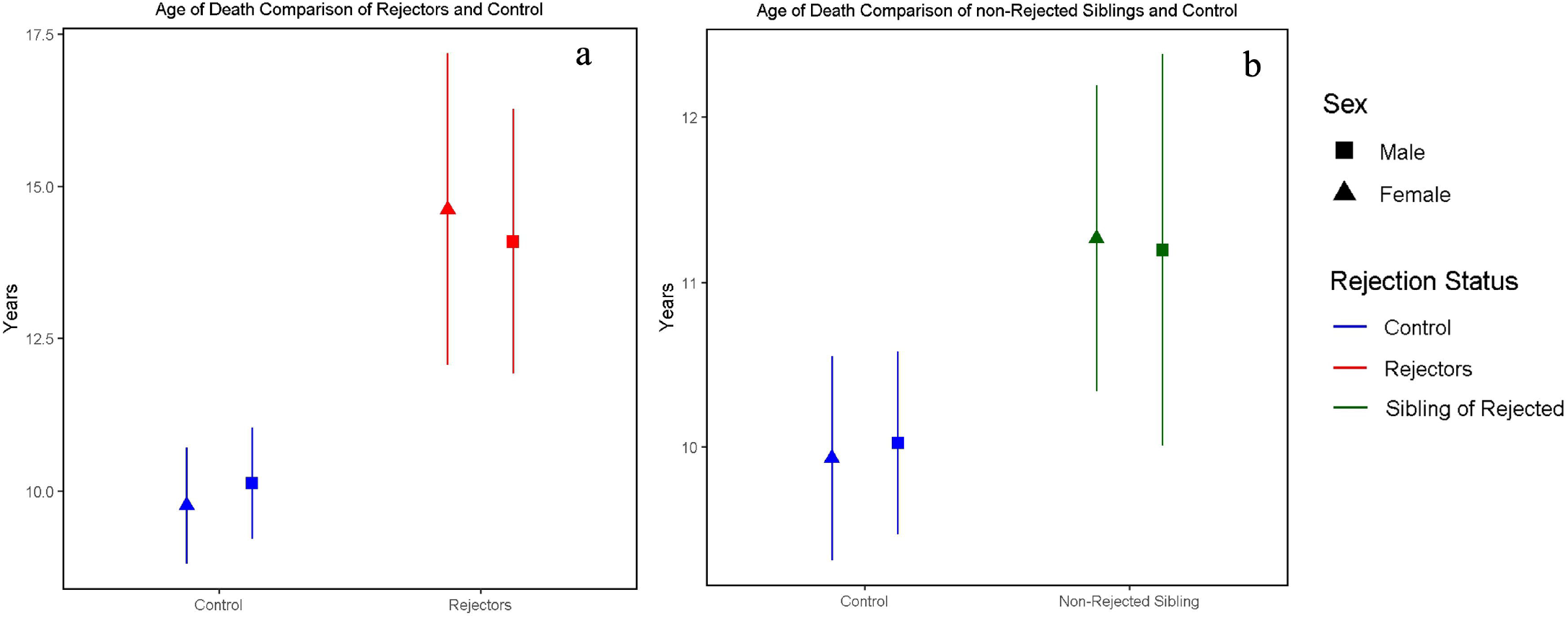
Lifespan effects of parental rejection in owl monkeys. (a) Rejector parents exhibit significantly longer lifespans compared to non-rejector parents. Median lifespan is indicated by symbols (triangles, rejector mothers n=19; non-rejector mothers n= l 12; squares, rejector fathers n=24, and non-rejector fathers n=124) Welch’s t-tests: ***p < 0.001. (b) Non-rejected offspring born to rejector parents show extended lifespans relative to offspring of non-rejector parents. Boxplots display median lifespan and interquartile ranges for offspring of rejectors (n=231) and offspring of non-rejectors (n=524). General Linear Model: p = 0.003.

### Offspring survival analyses

To assess whether having a rejecting parent conferred longevity benefits to well-reared offspring, we compared non-rejected siblings from rejector families (i.e., offspring raised by parents who rejected at least one sibling; n= 231) to non-rejected offspring from non-rejector families (i.e., offspring whose parents never exhibited rejection behaviour; n= 524). A generalised linear model was used with parental rejection status as the main predictor and sex as a covariate (Figure 1b). Outcomes for rejected offspring who survived to adulthood have been reported previously^4^.

### Parent Survival distribution

To ascertain when during the lifespan survival differences occurred between rejector and non-rejector parents (described above), we conducted survival analyses using the survfit() function from the survival package in R. Kaplan–Meier curves and log-rank tests were used to visualise and compare survival distributions (Figure 2a).

**Figure 2.**
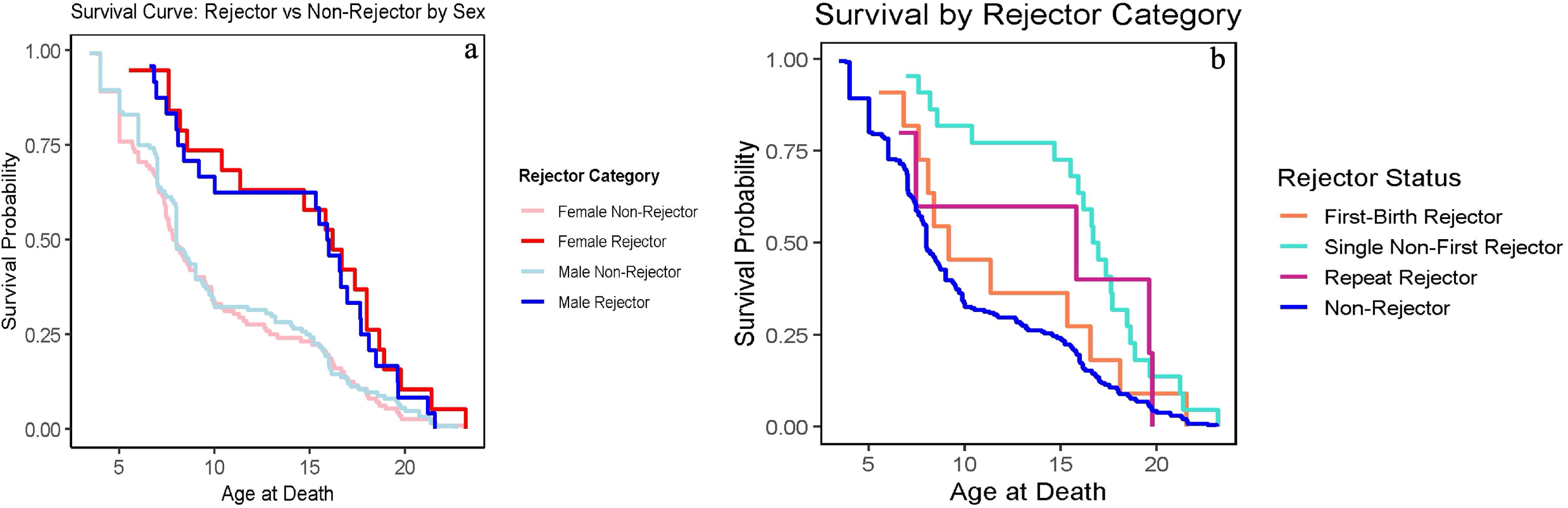
Kaplan-Meier survival curves comparing rejector and non-rejector owl monkey parents. (a) Survival probability by parental rejector status. Rejector mothers (red line, n=19) and rejector fathers (blue line, n=24) demonstrate enhanced survival compared to non-rejector mothers (pink line, n=l 12) and non-rejector fathers (light blue line, n=124), with the most pronounced differences during reproductive years. Log-rank test: [χ^2^ =14.8, df = 3, p =0.002]. (b) Survival probability by timing and frequency of rejecting behaviour. First-Birth Rejectors (n=l 1 ), Single Non-First Rejectors (n=22), and Repeat Rejectors (n=5) show distinct survival trajectories, with First-Birth Rejectors overlapping controls, while Single Non-First and Repeat Rejectors show visually elevated survival during middle adulthood. Log-rank test: [χ^2^ = 6.3, df = 3, p = 0.1].

For a subset of 38 individuals with detailed rejection histories, we further classified parents into three categories: repeat rejectors (n = 5, Male = 3 Female = 2), single non-first rejectors (n = 22, Male = 10, Female = 12), and first-birth rejectors (n = 11, Male = 8, Female = 3). This analysis was exploratory due to limited sample sizes. We first applied a General Linear Model (GLM) to test whether rejection category predicted lifespan. Similarly, to account for censored cases (individuals without a recorded age at death), we conducted survival analyses using the survfit() function from the survival package in R. Kaplan–Meier curves and log-rank tests were used to visualise and compare survival distributions (Figure 2b). Some rejector parents were excluded from these timing-specific analyses because the exact birth at which rejection occurred could not be determined, but they were retained in all broader lifespan and reproductive analyses.

To further explore differences in lifespan across rejector categories, we conducted a one-way ANOVA comparing age at death among the four original rejector groups (non-rejectors, first-birth rejectors, single non-first rejectors, and repeat rejectors), followed by post-hoc pairwise comparisons using Tukey’s HSD test. To complement this analysis and leverage larger group sizes, we also grouped individuals into later-birth/repeat rejectors (combining repeat rejectors and single non-first rejectors, n = 27; Male = 13, Female = 14) and comparison individuals (combining first-birth rejectors and non-rejectors, n = 11 + 236; Male = 132, Female = 115), and compared mean age at death between these groups using a Welch two-sample t-test.

### Reproductive success analyses

Reproductive success was measured as the total number of offspring produced. We analysed reproductive data for 279 individuals with known lifetime reproductive histories (summing to a total of 755 offspring), of which 43 exhibited rejection behaviour. Analyses of rejection timing or birth order necessarily used smaller subsets due to incomplete historical records; reproductive output analyses reported here use the full dataset of individuals with complete lifetime reproductive histories. A General Linear Model (GLM) was used to test the effects of rejector status and sex, as well as their interaction, on total reproductive output (Figure 3).

**Figure 3.**
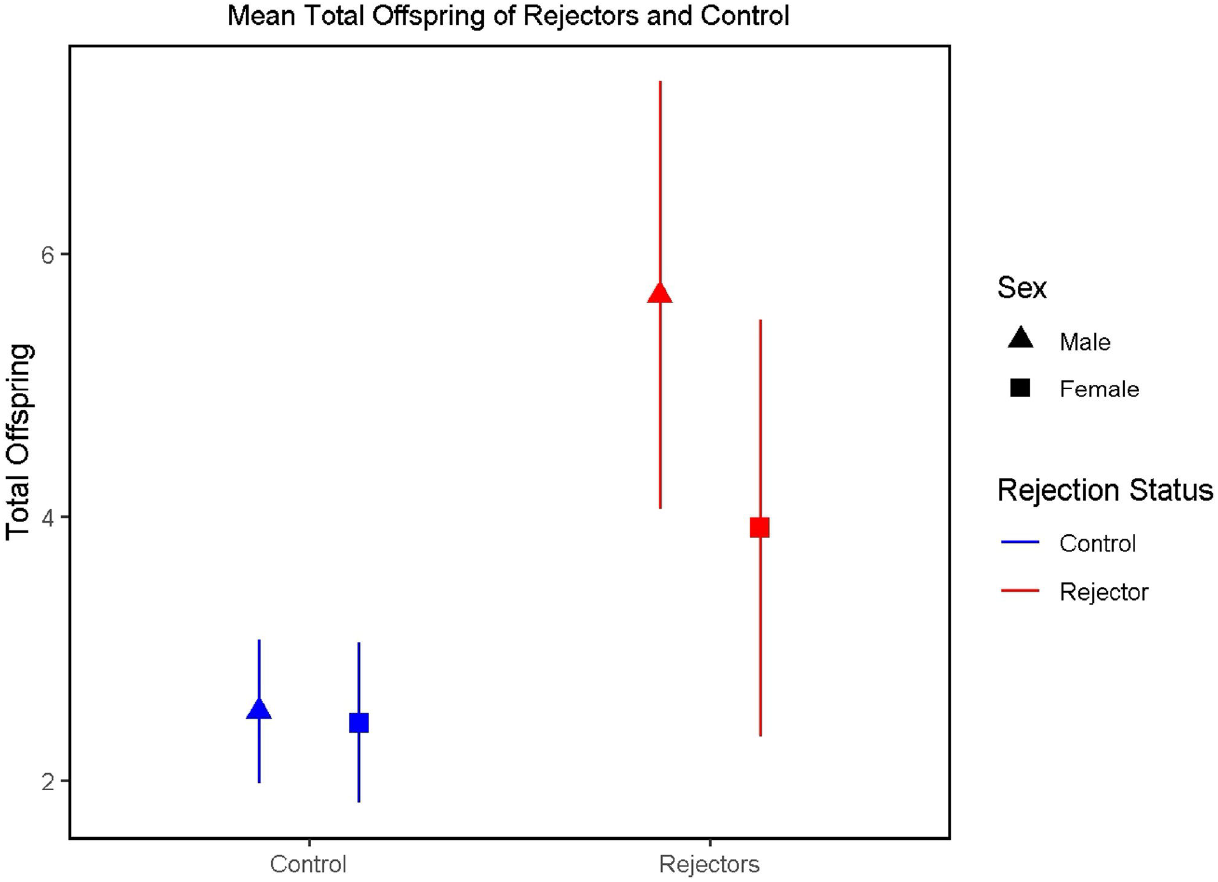
Reproductive output differs by rejector status and sex in owl monkeys. Total number of offspring produced by rejector versus non-rejector parents, shown separately for females and males. Points indicate median reproductive output. A General Linear Model revealed that rejector parents produced significantly more offspring than non-rejectors (p < 0.001). Parental sex did not significantly predict total reproductive output (p = 0.376).

### Statistical analyses

Welch’s t-tests were employed for lifespan comparisons between rejectors and controls. General Linear Models (GLMs) were fitted to test predictors of lifespan and reproductive output. Survival analyses were performed using the survival package in R, incorporating censoring where appropriate. All analyses were conducted in R (R Core Team, R 4.5.1).

## Results

### Longer lifespan expectancy in rejecting parents and their well-reared offspring

In owl monkeys, parental rejection of newborn offspring was associated with extended lifespan for both the parents and the offspring they raised. Comparing rejector parents with non-rejectors, Welch’s two-sample t-tests showed that rejector mothers (n = 19) lived significantly longer than non-rejector mothers (n = 112) (t = -3.17, p = 0.001, 95% CI [2.16, 7.57]), with mean ages at death of 14.33 and 9.95 years, respectively. Similarly, rejector fathers (n = 24) lived longer than non-rejector fathers (n = 124 (t = -3.46, p = 0.002, 95% CI [1.63, 6.31]), with mean ages at death of 14.16 years and 10.12 years, respectively (Figure 1a).

We also assessed whether well-reared offspring raised by rejecting parents benefited in terms of lifespan. These offspring (n = 231) lived on average 1.26 years longer than offspring from non-rejector families (n = 524) (coefficient = 1.26, SE = 0.43, t = 2.96, p = 0.003). Offspring sex (male vs. female) did not significantly affect lifespan (t = -0.19, p = 0.85) (Figure 1b). These results indicate that parental rejection is associated with longevity benefits both for the parents and for their well-reared offspring.

### Age-specific survival of rejector versus non-rejector parents

We used Kaplan-Meier survival curves to examine survival over the lifetime, to assess age-specific costs and benefits of rejecting behaviour, and to reveal how parental strategies influence longevity at different life stages. These curves included 19 rejector mothers, 112 non-rejector mothers, 24 rejector fathers, and 124 non-rejector fathers. Survival curves began to diverge early in life, with the greatest separation observed between ages 6-20 years. After ∼20 years, survival probabilities of males converged, and around 23 years for females. Censored individuals contributed information up to the age at last observation. The log-rank test confirmed that survival differences between rejector and non-rejector parents were statistically significant (χ^2^ = 14.8, df = 3, p = 0.002) (Figure 2a).

### Lifespan expectancy patterns associated with parental experience: effects of birth order on rejecting behaviour

To assess whether rejection timing differed between patterns commonly attributed to maternal inexperience and those occurring later in the reproductive lifespan, we compared survival trajectories of parents who rejected only their first offspring (First-Birth Rejectors, n = 11, Male = 8, Female = 3 ) with those who rejected at later births-either once (Single Non-First Rejectors, n = 22, Male = 10, Female = 12 ) or repeatedly (Repeat Rejectors, n = 5, Male = 3, Female = 2) versus non-rejectors (112 females, 124 males, n= 236). If first-birth rejection reflects inexperience, these individuals should not show enhanced longevity. A General Linear Model (GLM) tested whether rejection category predicted age at death while controlling for sex. Lifespan differed among rejection categories, with Single Non-First Rejectors living significantly longer than the reference category (First-Birth Rejectors). Single Non-First Rejectors showed an estimated 4.28-year increase in age at death (estimate = 4.28, SE = 1.89, *t* = 2.26, *p* = 0.025). In contrast, Repeat Rejectors did not differ significantly from First-Birth Rejectors (estimate = 2.23, SE = 2.76, *t* = 0.81, *p* = 0.419), nor did Non-Rejectors (estimate = −1.63, SE = 1.58, *t* = −1.03, *p* = 0.303). Sex was not a significant predictor of lifespan (estimate = 0.48, SE = 0.62, *t* = 0.77, *p* = 0.44). Overall model fit was moderate (AIC = 1678.5; residual deviance = 7026.3 on 269 df). Log-rank tests did not reach statistical significance (χ^2^ = 6.3, df = 3, p = 0.1) (Figure 2b). These results indicate that longevity benefits are most pronounced among parents engaging in single, non-first rejection.

To further examine lifespan variation across rejector categories, we conducted a one-way ANOVA comparing age at death among Non-Rejectors, First-Birth Rejectors, Single Non-First Rejectors, and Repeat Rejectors. The analysis revealed a significant overall effect of rejector category on age at death (F(3, 270) = 9.81, p < 0.001), indicating meaningful differences in lifespan across groups.

Post-hoc pairwise comparisons using Tukey’s HSD showed that Single Non-First Rejectors lived significantly longer than Non-Rejectors (mean difference = 5.88 years, 95% CI [2.94, 8.83], p < 0.001). No other pairwise contrasts reached statistical significance, including comparisons involving First-Birth Rejectors or Repeat Rejectors (all p > 0.12). Descriptively, Single Non-First Rejectors exhibited the highest mean age at death (mean = 15.8 years), followed by Repeat Rejectors (mean = 13.9), First-Birth Rejectors (mean = 11.7), and Non-Rejectors (mean = 9.95).

Although residuals deviated from normality (Shapiro–Wilk W = 0.92, p < 0.001), ANOVA is robust to such violations given the large sample size and relatively balanced variances. However, statistical power was limited for smaller categories, particularly Repeat Rejectors (n = 5) and First-Birth Rejectors (n = 11), which may have constrained detection of additional differences. Overall, these findings suggest that lifespan differences are most pronounced among parents engaging in single, non-first rejection, rather than rejection per se.

To complement this analysis, we grouped individuals into later-birth/repeat rejectors (Repeat Rejectors and Single Non-First Rejectors, n = 27) and comparison individuals (First-Birth Rejectors, n = 11 plus Non-Rejectors, n = 236), and compared their mean age at death using a Welch two-sample t-test. Later-birth/repeat rejectors showed significantly higher mean age at death compared to comparison individuals (15.47 vs. 10.03 years; t(32.40) = -5.40, p < 0.001, 95% CI = -7.49 to -3.39). This result suggests that individuals exhibiting rejection beyond the first birth tend to live longer than first-birth rejectors and non-rejectors. The t-test complements the ANOVA by leveraging larger group sizes for these grouped classifications, providing additional statistical power. Sex-specific analyses revealed that this lifespan difference was present in both females and males. Among females, later-birth/repeat rejectors lived significantly longer than comparison females (15.53 vs. 9.72 years; t(16.35) = -4.07, p < 0.001, 95% CI = -8.83 to -2.79). Similarly, later-birth/repeat male rejectors showed significantly greater longevity than comparison males (15.40 vs. 10.30 years; t(14.61) = -3.47, p = 0.003, 95% CI = -8.25 to -1.96).

Together, these results indicate that individuals exhibiting rejection beyond the first birth, or across multiple offspring, consistently achieve longer lifespans, regardless of sex. This pattern differs from first-birth rejection, which is commonly attributed to parental inexperience. By aggregating rejection categories into later-birth/repeat rejectors and comparison individuals, this analysis increases statistical power and reinforces the patterns observed in the rejector-category ANOVA.

### Reproductive output of rejector parents is greater than for controls

We assessed whether rejector parents produced more offspring than non-rejectors and whether this effect varied by sex. A General Linear Model (GLM) tested the effects of rejector status (rejector vs. non-rejector), parental sex (male vs. female), and their interaction on total reproductive output.

Rejector parents produced significantly more offspring than non-rejectors (estimate = 2.23, SE = 0.54, *t* = 4.12, *p* < 0.001). Sex did not significantly predict offspring number (estimate = −0.35, SE = 0.39, *t* = −0.89, *p* = 0.376), and the interaction between rejector status and sex was not significant (estimate = −1.68, SE = 1.09, *t* = −1.54, *p* = 0.124), indicating that the reproductive advantage of rejection was similar in males and females.

Because lifespan constrains total reproductive output, we further tested whether rejector status predicted total offspring after accounting for age at death. Lifespan was a strong predictor of reproductive output (estimate = 0.46, SE = 0.03, *t* = 15.49, *p* < 0.001), whereas rejector status was no longer significant (estimate = 1.16, SE = 1.11, *t* = 1.05, *p* = 0.295). The interaction between rejector status and lifespan was also non-significant (estimate = −0.066, SE = 0.075, *t* = −0.88, *p* = 0.379). These results indicate that the previously observed reproductive advantage of rejectors is largely explained by their longer lifespan rather than by increased reproductive output per year.

Overall, rejector parents produced more offspring, but this effect is fully accounted for by longevity, with no evidence that males and females differed in the reproductive benefit of rejection.

A Welch two-sample t-test comparing total offspring between rejectors and non-rejectors indicated that rejectors produced significantly more offspring overall. Across the full sample, rejectors (n = 43) had a mean of 4.70 offspring, compared with 2.48 for non-rejectors (n = 236), t(54.28) = –3.73, *p* < 0.001, 95% CI [–3.41, –1.03].

Sex-stratified analyses revealed that this effect was significant in females but not in males. Female rejectors (n = 19) produced an average of 5.68 offspring, compared with 2.53 for non-rejecting mothers (n = 112), t(22.79) = –3.85, *p* < 0.001, 95% CI [–4.85, –1.46]. Among males, rejectors (n = 24) had a mean of 3.92 offspring, compared with 2.44 for non-rejecting fathers (n = 124), t(30.91) = –1.79, *p* = 0.083, 95% CI [–3.17, 0.20], indicating no significant difference.

These results suggest that parental rejection is associated with higher lifetime reproductive output overall. Sex stratified analyses indicate this difference is statistically robust in females, while male rejectors do not significantly differ from non-rejectors in offspring number.

## Discussion

We explored the biological significance of infant rejection in a captive colony of owl monkeys. Parents who rejected their offspring showed considerably longer survival and higher reproductive rates than those who did not, and these benefits extended to their offspring whom they successfully raised. These findings are consistent with the hypothesis that reduction in parental investment through rejection may reflect altered allocation of care away from specific offspring^31^.

Although these analyses are based on a captive colony, our study enables the study of rare and extreme parental behaviours under longitudinal conditions rarely achievable in the wild. Neonatal abandonment and severe rejection are not artefacts of captivity, but captivity may increase their frequency by amplifying ecologically relevant stressors. Rather than generating novel behaviours, captive conditions likely shift individuals along an existing continuum of parental investment strategies, exposing trade-offs between offspring care, parental condition, and future reproduction.

### Survival in rejecting parents and offspring

Parental rejection was associated with increased survival in both mothers and fathers. Rejector mothers lived longer than non-rejecting mothers (14.33 vs. 9.95 years), and rejector fathers lived longer than non-rejecting fathers (14.16 vs. 10.12 years) (Figure 1a). Both sexes showed similar patterns, which is consistent with the biparental care system of owl monkeys, where both parents assume a considerable energy cost in raising the young. Sex-specific parental strategies may contribute to explaining survival differences. In females, extended survival may reflect the conservation of resources that would otherwise be used for lactation, an energetically costly process^44^ that continues in owl monkeys until weaning at 7-8 months^45^. In males, the absence of infant carrying duties could provide equivalent energy savings.

We also demonstrated that well-reared offspring from rejecting families survived significantly longer than offspring from non-rejecting families: 1.26 years longer on average (Figure 1b). These gains were intermediate between those of rejecting parents (≈4-4.5 years longer lifespan) and controls, suggesting that parental phenotype, rearing experience and the behavioural act of rejection may contribute to different outcomes. In offspring, a possible explanation for the longer life expectancy may reflect having a rejector as an ancestor. But unlike their rejected siblings, well-reared offspring enjoy secure attachment in early life and are breastfed and cared for by parents with lower energetic demands than controls, which has been shown to confer advantages in macaques^46^.

The increase of average life expectancy was primarily due to increased survival during the reproductive years - between ages 6 - 20 years (which can extend to 24 years in this colony^46^)-rather than a prolongation of maximum lifespan. Survival curves converged ∼ 21 years, and both parents who rejected newborns and those who did not showed similar mortality rates in extreme old age (23+ years, Figure 2a). This pattern indicates that rejection is associated with improved survival during the energy-intensive reproductive period but does not alter intrinsic ageing processes.

Our study of survival patterns depending on the occurrence of rejection by birth order showed no statistically significant effects. This study was underpowered, however, consistent visual differences in Kaplan-Meier survival trajectories were observed. Rejection in the first birth -widely documented in captive primates and attributed to maternal inexperience-showed no survival advantage, with trajectories overlapping with those of the controls (Figure 2b). In contrast, individuals rejecting offspring beyond the first birth maintained substantially higher survival probabilities starting from 8 years, and this advantage becomes more pronounced in later life (10-15+ years). Repeat rejectors showed similarly elevated survival during middle adulthood. Exploratory ANOVA and t-testing support these visual trends.

Individuals rejecting offspring after their first birth (either once or repeatedly) lived significantly longer than those that did not exhibit this pattern, with a mean difference of approximately 4 years (median ∼16 years). This finding is consistent with the possibility that rejection occurring after the first birth reflects responses to resource constraints or environmental conditions, rather than maternal inexperience. Longer lifespan appears confined to individuals who reject later in their reproductive lives and possess experience rearing offspring. Such rejection in later life stages may reflect differences in resource allocation under energetic stress, consistent with observations in wild baboons, where rejecting rates vary with environmental conditions^11^. Reaching old age can be characterised by worsening health in non-human primates^48^, and behavioural disorders such as rejecting could be accompanied by comorbidities^49^. Future studies will characterise healthspan and gerospan in this long-lived rejector cohort and their well-reared descent.

### Reproductive performance and the compensation challenge

Parents who rejected their offspring produced significantly more offspring than those who did not: males had an average of 3.92 offspring compared to 2.44, and females an average of 5.68 compared to 2.53 (Figure 3). This research revealed a nuanced picture of disposable soma theory in owl monkeys in this colony. In *Aotus* spp., where both parents share caregiving^43^, rare among mammals, the theory’s predictions held: both sexes showed similar longevity gains as expected. However, parents who rejected offspring do not fit simple expectations of the theory: they lived longer and produced more total offspring, by virtue of living longer. The expected trade-off, which is typically obscured in natural populations but measurable under controlled conditions^50^, was not observed here. Rejector pairs also reproduce less frequently, as even though life expectancy is substantially extended, they only have statistically one or two more offspring. In this colony individuals can continue reproduction beyond 20 years^47^, then it is unknown if fertility in mother rejectors might be impaired.

When rejecting parents face energy shortages that threaten their survival, withdrawal of care from offspring may be associated with reduced physiological costs and improved survival, potentially increasing total lifetime reproductive output^6,11,36,51^. Rather than directly increasing reproductive rate, rejection and slower reproduction may influence the timing and allocation of resources throughout life, affecting the balance between reproduction and somatic maintenance under stressful conditions. This mechanism may be particularly relevant in owl monkeys, which are vulnerable to stress^52^. In humans, stress increases cellular energy expenditure by 60–110%^34^. Elevated stress can constrain reproductive capacity, reducing reproductive output even in otherwise healthy captive non-human primates^53^. Under these conditions, the energetic costs of reproduction might overwhelm owl monkey mothers with heightened stress physiology. Redirection of energetic investment from reproduction to somatic maintenance may contribute to extended survival of rejectors and their well-reared offspring.

As only a few individuals become rejectors whilst living under the same conditions, differences in life expectancy likely arise from variation in genetic background, experience of parental investment, and the behavioural act of rejecting itself. If longer life expectancy were purely genetic, rejector parents and all their descendants would have similar lifespans; if purely behavioural, only individuals who themselves reject would survive longer; and if purely environmental, more individuals would reject and well-reared offspring of rejectors would not survive longer than well-reared controls.

Molecular mechanisms underlying these patterns remain to be determined. Molecular factors associated with stress and longevity have been observed in some rejectors’ well-reared offspring. As observed previously, rejecting is more commonly observed in individuals with genetic variants associated with environmental sensitivity and serotonergic neurotransmission^28,29^. These variants are susceptible to epigenetic modifications, induced by environmental stress and contribute to manifestation of altered behavioural states in adulthood. In owl monkeys, overexpression of miR-125b-5p (and Let7c) is detectable in well-reared offspring of rejectors in infancy (∼1 year), before they reproduce^5^. These biomarkers were also identified in humans exposed to early life stress that later develop stress triggered behavioural disorders, with authors demonstrating epigenetic reprogramming of genetic variants in the miR-125b loci created gene expression differences expressed in the brain^54^. miR125-5p together with Let7 also regulate key longevity pathways, such as Chinmo55, p^53,56^, and are neuro and cardioprotective biomarkers^57,58^ and tested in cell regeneration studies^59^. Excessive stress and cardiomyopathies are significant causes of death in this and other colonies of owl monkeys^60,61^, thus if proven functional, observed miRNA profiles could be protective. However, these molecular signatures were observed in rejectors exposed to early life rejection, not the entire sample, which is heterogeneous. Multiple pathways likely contribute to its manifestation.

Additionally, we previously showed parenting experience is associated with accelerated epigenomic ageing affecting brain rather than blood tissue in this colony^3^. Further, some rejecting parents are characterized by biomarkers associated in humans with non-alcoholic fatty liver disease and present agalactia, a condition linked to inflammatory and metabolic alterations^5,62^. Reduction in parental investment and rejection occur in response to interventions which lower fat content in diet in non-human primates^63^. These observations raise the possibility that, in absence of noticeable weight differences, might occur in response to reduction of specific lipid content and disrupt nursing before reaching critical exhaustion of reserves destined to somatic repair of organs like the heart and the brain. In future studies we will test if rejectors possess allelic variation associated with greater stress sensitivity, display altered fat metabolism and enjoy slow epigenetic ageing.

The scientific literature consistently describes rejection as a pathological behaviour that occurs under stress, poor health, and compromised well-being in humans and non-human primates^64-66^. Experimental maternal separation in non-human primates is used as a valuable model for human psychiatric conditions and their health repercussions stemming from insecure attachment^65-69^. The convergence of molecular markers we identified in owl monkeys^4,5^ and others in humans^54,70^ in individuals exposed to early life stress that develop behavioural and neuropsychiatric disorders speak of evolutionary conserved mechanisms linking exposure and adult behavioural pathology.

These results suggest that behaviours associated with early-life stress may have measurable consequences for survival and health. Rather than being solely pathological, such behaviours may also reflect responses to environmental or physiological constraints. Whether these responses represent conserved mechanisms across species remains an open question.

Despite being associated with longer lifespan, the rejector phenotype remains infrequent (5-10%) in this colony and stable across generations, suggesting a balance between potential survival benefits and the costs associated with offspring mortality. While also observed in other captive Aotus colonies and in many other captive primate colonies, whether comparable extension in life expectancy arises where rejecting is reported remains to be determined.

As some rejected individuals that become rejectors are also characterised by longer than average life expectancy^5^, it is possible fitness in rejected-rejectors, and their offspring could also be enhanced. In future, we will study this phenomenon as it might constitute an example of a nongenetic mechanism that facilitates rapid adaptation to chronic environmental challenges, such as captivity, in timescales not amenable to genetic evolution.^74,75^

## Conclusion

Our findings show that parental rejection is associated with extended lifespan and higher lifetime reproductive output in a captive population of owl monkeys. These patterns highlight the importance of considering variation in parental investment when interpreting life-history trade-offs under constrained conditions. While rejection is typically viewed as pathological, our results suggest that it may also reflect shifts in resource allocation with measurable consequences for survival. Determining whether similar patterns occur in natural populations will be important for understanding the broader biological significance of this behaviour.

## Ethics statement

All data analysed in this study derive from long-term veterinary, demographic, and behavioural records collected as part of routine colony management at the Centre for Reproduction and Conservation of Non-Human Primates (CRCP), IVITA, Universidad Nacional Mayor de San Marcos (UNMSM), Peru. No experimental procedures or interventions were performed specifically for this study. Animal husbandry and veterinary care complied with the American Society of Primatologists (ASP) Principles for the Ethical Treatment of Non-Human Primates. Oversight and approval were provided by the Welfare and Research Committee of San Marcos University and the UNMSM ethics committee. Research activities at the centre were conducted under permits issued by the Peruvian authority SERFOR to Dr. Paredes (AUT-IFS-2021-040 and RDG No. D000334-2021-MIDAGRI-SERFOR-DGGSPFFS).

## Data availability

The demographic, behavioural, and reproductive data analysed in this study derive from long-term colony management records maintained at the Centre for Reproduction and Conservation of Non-Human Primates (CRCP), IVITA, Universidad Nacional Mayor de San Marcos (UNMSM), Peru. Due to ethical, legal, and institutional restrictions governing access to non-human primate records, the raw datasets are not publicly available. De-identified data supporting the findings of this study are available from the corresponding author upon reasonable request and subject to approval by CRCP-IVITA and UNMSM.

## Author Contributions

**JF:** formal analysis (led), investigation (equal), draft revision (equal). **NS:** sampling (led), resources (equal) draft revision (equal). **PY** draft revision (equal). **UMP:** conceptualization (lead), investigation (equal), original draft (lead).

## Acknowledgements

We thank the technical and medical staff at the IVITA centre for Primate Conservation and Breeding, UNMSM, for their assistance with sample collection and compilation of data. UMP would like to thank Dr. Riadh Abed from the Evolutionary Psychiatry Special Interest Group (EPSIG), Prof. Sarah Blaffer-Hrdy and Dr. Guy Cowlishaw and the UCL Biological Anthropology community for helpful discussions.

